# HMGB1 mediates macrophage recruitment and regional intervertebral disc tissue functional and mechanical property changes following injury

**DOI:** 10.1101/2024.10.30.621107

**Authors:** Kevin G. Burt, Min Kyu M. Kim, Dan C. Viola, Joseph R. Genualdi, Gerard F. Marciano, Nadeen O. Chahine

## Abstract

**Objective:** Frequently evaluated in musculoskeletal disease, damage associated molecular patterns (DAMPs) respond to tissue damage and cellular stress by facilitating an inflammatory response via macrophage activation and broad inflammatory pathway activation. In the context of disc degeneration (DD), high mobility group box 1 (HMGB1), a potent intracellular DAMP, is seen to be increased within severely degenerated human IVDs and to directly mediate inflammatory responses within disc cells *in vitro*. To further understand how HMGB1 mediated inflammation influences DD, this study evaluated the possible protective effect of an HMGB1 knockout on DD pathology following injury.

**Methods:** Using a needle puncture injury model in murine caudal IVDs we evaluated DD pathology within an IVD specific *Hmgb1* knockout (KO) model. Structural and compositional changes in IVD cellularity, histopathology, disc height, and biomechanics were evaluated in addition to an assessment of disc inflammation via gene expression and macrophage presence throughout the course of degeneration.

**Results:** HMGB1 expression robustly increased shortly following needle puncture injury and elevated levels were sustained up to 28-days post injury both in injured IVDs and in the IVDs adjacent to the level of injury. IVD specific *Hmgb1* KO mice had an increased disc height following injury both at the injured and adjacent to injury level compared to injured WT IVDs. Hmgb1 KO also protected against tissue mechanical property losses at both the injured (dynamic modulus) and adjacent to injury level (dynamic modulus, creep, and equilibrium modulus) compared to injured WT IVDs, however there was no significant effect on histopathologic scores post injury. *Hmgb1* KO resulted in alterations in macrophage (F4/80+) recruitment to the IVD post injury *in vivo.* A lower macrophage migration was also observed *in vitro* in response to the secretome of an injured *Hmgb1* KO IVD compared to injured WT IVDs. *Hmgb1* KO had no effect on inflammatory gene expression changes following injury within adjacent to injury level or injury level IVDs.

**Conclusion:** Overall findings indicate that HMGB1 is upregulated regionally, at both the injured level and at the level adjacent to injury. Results suggest that HMGB1 plays a role in mediating structural, biomechanical, and inflammatory responses to IVD injury and serves as a potent chemoattractant, mediating macrophage recruitment to the IVD and overall migratory function.

## Introduction

With degeneration, the intervertebral disc (IVD) undergoes structural and biological changes producing a mechanically compromised, chronic inflammatory, and catabolic microenvironment. Structural hallmarks of disc degeneration (DD) include a loss of cellularity, ECM components, and cartilaginous structure, and overall loss of disc height [1–6]. Biological hallmarks of DD include chronic upregulation of inflammatory cytokines, including: TNFα, IL-1β, IL-6, and IL-17 [7–11], catabolic enzymes (MMPs and ADAMTS) [12–14], and increased presence of macrophages, a phagocytic innate immune cell [15–17]. In addition to these inflammatory mediators, damage associated molecular patterns (DAMPs) have also been associated in the mediation of chronic inflammation during DD.

DAMPs are molecules released from cells and tissues in response to damage, most often injury at the tissue level, that result in stress at the cellular level. DAMPs were originally proposed as an an immune response activated not by the signals previously defined as “self” vs. “non-self”, but rather as a detection and protection from danger [18]. Since then, DAMPs originating from both the extracellular and intracellular space have been identified following tissue injury or cell death. Following release DAMPs are initiate inflammatory responses via several different mechanisms including macrophage activation and downstream activation of pattern recognition receptors (PRRs) [19]. Extracellular DAMPs, released following tissue damage, include ECM components, biglycan, fibronectin, and hyaluronan [20]. Intracellular DAMPs, released following cell stress or death, include histones, S100 proteins, heat-shock proteins (HSPs), and HMGB1 [20]. Within musculoskeletal tissues, chronic inflammation, cell death, and wide scale tissue degeneration are common hallmarks of disease. As such, DAMPs are often evaluated for their roles in responding to musculoskeletal tissue damage throughout the onset of degeneration, and during chronic inflammation. Specifically, HMGB1 has been shown to play a role in many musculoskeletal diseases including tendinopathy, osteoarthritis (OA), rheumatoid arthritis (RA), and DD [21–27].

With substantial evidence supporting HMGB1 upregulation in human samples with musculoskeletal diseases, studies have evaluated the role of HMGB1 within animal models of bone fracture, muscle injury, and arthritis [28–35]. Fracture and muscle injury models revealed a beneficial effect of exogenous HMGB1 treatment, where better healing outcomes were observed based on increased bone formation at fracture site, increased bone strength, and an earlier increase in recruitment of circulating leukocytes and neutrophils when compared to the vehicle treated controls [28]. Alternatively, animal models of arthritis have observed that treatment with exogenous HMGB1 leads to worsened outcomes including histological tissue degradation and swelling within joints [29–32]. Whereas inhibition of HMGB1 within such arthritic models led to a mitigation of joint swelling, degeneration, and inflammation [29, 33–35].

Specific to the IVD, HMGB1 levels are higher in severely degenerated human IVDs compared to less degenerated tissues [22, 23]. Interestingly, *in vitro* treatment of human IVD cells with HMGB1 has produced a mixed response, with multiple studies observing an increase of pro-inflammatory cytokines (PGE_2_, TNF-α, IL-6, IL-1β) and catabolic enzymes (MMP-1, -3, and -9) [23, 36], while another found a minor inhibition of the pro-inflammatory cytokine, IL-6 [37]. Despite the evidence for upregulation of HMGB1 within degenerated human IVDs and the *in vitro* evaluations suggesting HMGB1 may contribute to DD through inflammatory activation of IVD cells, the role of HMGB1 in response to IVD injury *in vivo* has not been evaluated.

Studies using animal models to evaluate DD via induced structural damage have employed IVD puncture methods, where various gauges of needle are used to produce injury, typically herniation, in lumbar or caudal IVDs [38–44]. Such models have been developed in small rodents, such as rat [42] and mouse [38–41], in addition to larger animal studies including rabbit [43] and pigs [44]. A large benefit of utilizing mouse models is the ability to evaluate specific molecular mechanisms made possible by ease of genetic manipulation. Within these murine injury models, reproducible histological and ECM related changes have been observed alongside temporal upregulation of catabolic enzyme and inflammatory chemokine expression [38, 40]. However, the response of potent DAMPs, such as HMGB1, have not yet been explicitly studied within an IVD injury model.

The goal of this study was to evaluate a possible protective role of HMGB1 following injury to the IVD. To achieve this, we utilized an IVD specific *Hmgb1* knockout mouse and evaluated injury and healing responses following a caudal needle puncture injury. We hypothesized that a knockout of *Hmgb1* within IVD cells would lead to an attenuation of inflammatory gene upregulation and macrophage recruitment, overall leading to increased IVD healing following injury with maintenance of tissue composition and mechanical properties. Contrary to our hypothesis, we observed minimal effects of HMGB1 KO on IVD structural integrity, as indicated by histological analysis, or inflammatory gene expression in HMGB1^KO^ at the injured level or at the adjacent to injury. However, results suggested HMGB1 mediated changes to mechanical properties and disc height within injured levels and at the level adjacent to injury. Lastly, we observed macrophage recruitment *in vivo* and migration *in vitro* to be mediated by HMGB1 within injured IVDs. Overall, further providing support for the role of HMGB1 as a potent chemokine following disc injury, and the possible tissue mechanical and functional consequences downstream of this inflammatory role.

## Materials & Methods

### Mice

All mice used in the study were derived from C57BL/6J strain. Mice were housed in conventional cages and maintained on a 12-hour light/dark cycle, with rodent chow and water available *ab libitum*. At specified time-points, mice were euthanized by CO_2_ asphyxiation. Animal studies were conducted with approval by the Institute of Animal Care and Use Committee of Columbia University (Protocol #: AC-AABQ8555).

### HMGB1 knockout Mice

Conditional *Hmgb1* knockout mice (*HMGB1 ^fl/fl^,* JAX stock no. 031274) containing loxP sites flanking exons 2 through 4 of the *Hmgb1* gene were used to target the deletion of HMGB1 [45]. Homozygous *HMGB1 ^fl/fl^* mice were bred to mice heterozygous for aggrecan (*Acan*) knock-in allele carrying tamoxifen-inducible form of Cre recombinase (*Acan^CreERT2/+^*; JAX stock no. 019148) [46]. Mice without CreER^T2^ recombinase were used as controls (*Acan^+/+^;HMGB1^fl/fl^*, hereinafter referred to as Control) for comparison to CreER^T2^-positive mice (*AcanCre^ERT2/+^;HMGB1^fl/fl^*, hereinafter referred to as HMGB1^KO^). Cre-mediated recombination, resulting in HMGB1 deletion, was induced in skeletally mature (3-4 months-of-age) mice via intraperitoneal tamoxifen injections (0.3 mg/g of body weight dissolved in sunflower seed oil; Sigma-Aldrich, Cat. No. T5648) for 3 consecutive days. Littermate control mice received the same tamoxifen injection. Following tamoxifen-induction, for baseline (uninjured) analysis control and HMGB1^KO^ mice were euthanized at 1-, 2-, and 3-months post-injection. Lumbar and caudal IVDs were harvested.

### Injury

A needle puncture injury was induced in caudal spine segments of skeletally mature (3-4 month old) mice. Prior to surgery an analgesic (Buprenorphine SR, 1.0 mg/kg) was administered via intraperitoneal injection. Mice were anesthetized by inhalation of Isoflurane (2-5%). Under anesthesia Co5/6 and Co7/8 IVDs were punctured percutaneously with a 26G needle inserted using fluoroscopic guidance (Glenbrook Technologies) until the needle reached two-third of disc width (Fig. 1A) [38]. Once inserted, the 26G needle remained within the IVD for 30s before 180 degree rotation and removal. Mice were euthanized for analysis at 1-, 3-, 7-, 14-, 28- and 56-days following the injury. An injured (Co5/6) and an adjacent uninjured disc (Co6/7) were snap-frozen for gene expression analysis (N=6 per level and time point). Another injured disc (Co7/8) and an adjacent uninjured disc (Co8/9) were fixed and decalcified for histological and immunohistochemically analysis (N=6 per level and time point). Age matched uninjured mice were used to isolate controls tissues for a basal comparison group (N=6). For HMGB1 KO studies, prior to injury HMGB1^KO^ and control mice received tamoxifen IP injections as described previously. Injury was induced within HMGB1^KO^ and control mice 3-days following IP injections and Cre-mediated recombination. Age matched uninjured HMGB1^KO^ and control mice were used to isolate IVDs for a basal comparison group (N=3-6).

**Figure 1:**
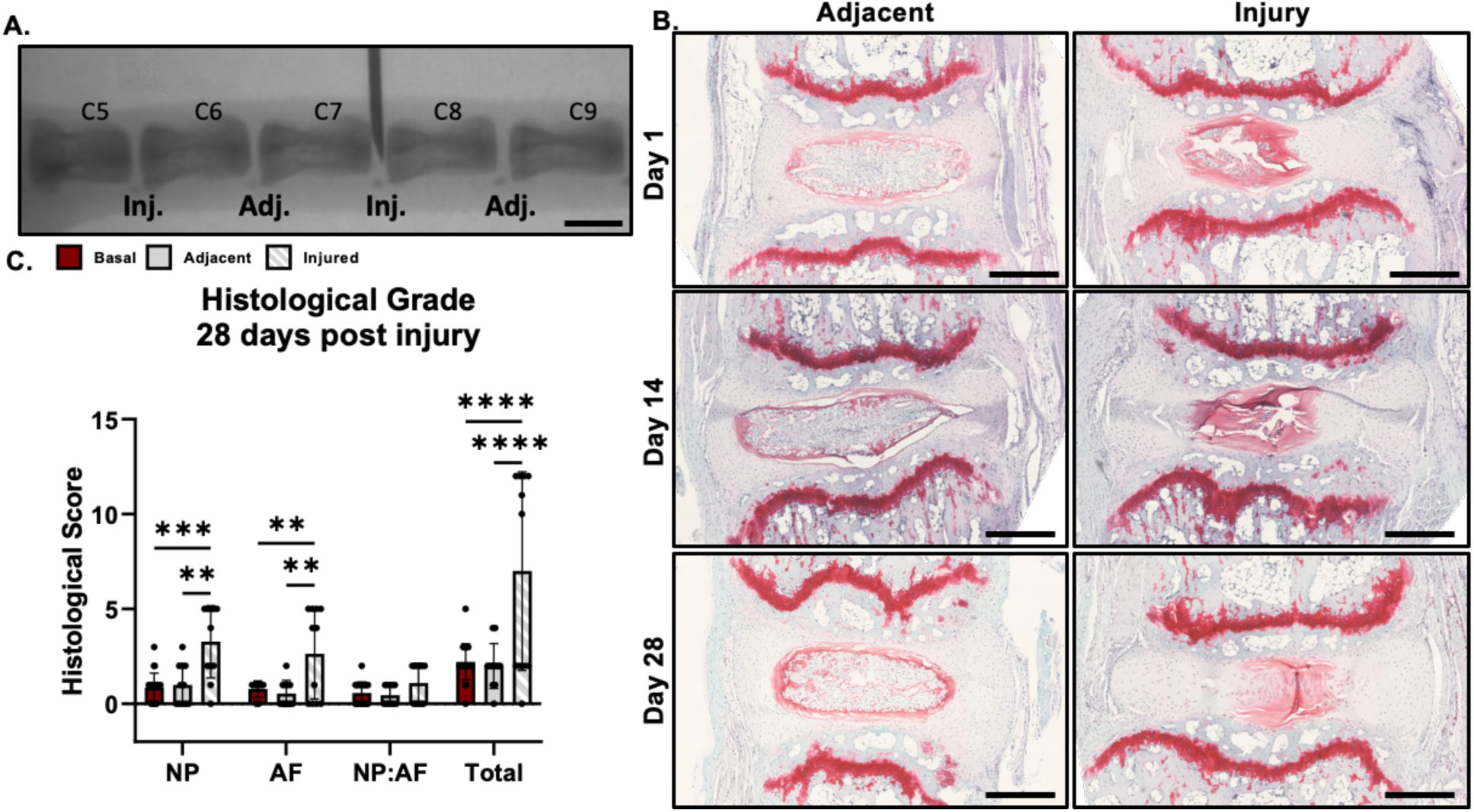
Validation of histological degeneration following caudal needle puncture injury. (**A**) Fluoroscopy image of needle puncture in the caudal spine. Scale bar = 1mm. (**B**) Representative Safranin-O stained mid-sagittal histological sections of injured and adjacent IVDs at 1-, 14-, and 28-days following injury. Scale bar – 500μm. (**C**) Histological scoring of 28-day injured, adjacent, and age matched uninjured (basal) IVDs. Histological scoring across NP, AF, NP:AF (border), and Total (summed score) compartments. **p<0.01, ***p<0.001, ****p<0.0001.

### Gene Expression

Whole disc tissues containing the NP, AF, and EP were isolated, snap frozen, and homogenized using a bead tissue homogenizer (Mikro-Dismembrator U, Sartorius) (N=6 per group). Total RNA extraction was performed using TRIzol (Thermofisher) and chloroform phase separation with RNA cleanup via spin columns (Qiagen) according to the manufacturer’s protocol. RNA purity and quantity was analyzed using a Nanodrop spectrophotometer (Thermofisher). For compartmental RNA isolation, NP and AF tissue was separated via microdissection, total RNA was extracted as described above with 8 levels (C4-C12) being pooled per mouse for one biological replicate (N=3-5). Complementary DNA (cDNA) was reverse transcribed using iScript cDNA synthesis kit (Bio-rad) and real time PCR was carried out with Itaq Universal SYBR kit (Bio-rad) and Applied Biosystems QuantStudio 7 Flex thermocycler. PCR thermocycler program included a 40 cycle cDNA amplification alternating between denaturation at 95°C for 15 sec and annealing/extension and plate read at 60°C for 60 sec. Relative gene expression was quantified using ddCT analysis for inflammatory cytokines (Table 1). Gene expression values were normalized to glyceraldehyde-3-phosphate dehydrogenase (*Gapdh*).

**Table 1:**
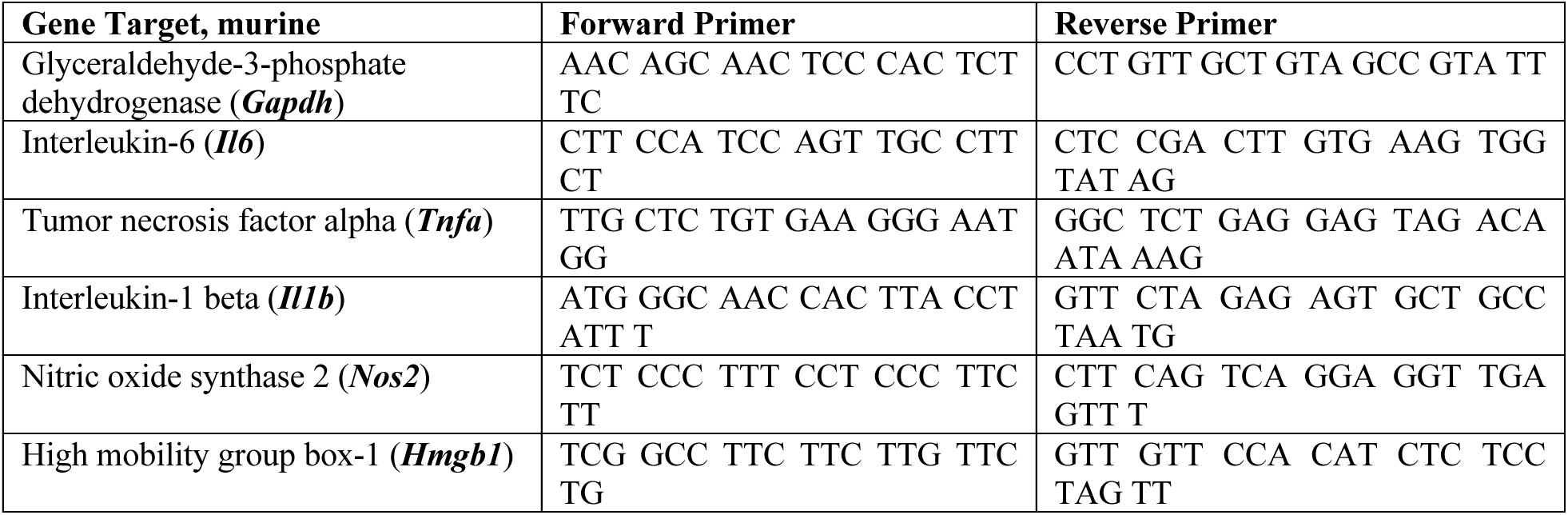
Forward and reverse gene primers sequences.

### Histological analysis

Bone-disc-bone spine segments (C5-C9; 4 IVDs/ animal/ genotype/ time point) were isolated, fixed in 4% paraformaldehyde for 24 hr, decalcified in 14% EDTA for 10 days, and either processed for paraffin-embedding or soaked in sucrose before cryo-embedding in OCT. Tissue structure was analyzed using paraffin embedded sagittal sections (7μm) stained with Safranin-O (cartilage/mucin)/Fast Green/Haematoxylin. Stained slides were imaged using an Axio Observer (Zeiss) using 10x/0.25 A-Plan objective, Axiocam 503 color camera, and Zen software.

Histomorphological analysis was performed using a mouse specific histological grading system, with higher scores indicating increased tissue degeneration [47]. IVD tissue processing was performed as described. Scoring of NP structure (scored 0-4) evaluated NP cellularity and clustering, with increased presence of “honey-comb” cell clustering, a loss of cellularity, and presence of mineralized matrix yielding higher scores. AF structure (scored 0-4) evaluated degenerative changes such as concentric lamellae alignment, widening or serpentine lamellae, rounded cell morphology, complete loss of structure, and mineralization. NP (scored 0-2) and AF (scored 0-2) clefts were evaluated for length in relation to compartment size. The NP/AF boundary health (scored 0-2) was assessed for a healthy defined compartment boundary and for degenerative changes consisting of rounded chondrocyte-like cells and discontinuity or complete loss of a clear boundary. Scoring was broken up into compartments including NP structure and clefts (0-6), AF structure and clefts (0-6), and NP/AF boundary (0-2), or combined for a total histological score (0-14). Stained sections were scored blinded to experimental groups. Scoring was analyzed comparing multi-level pooled discs between genotype groups.

### Immunofluorescence microscopy and image analysis

Paraffin embedded tissue sections were baked at 60°C for 35 min, deparaffinized with xylene, and rehydrated using graded series of ethanol washes. Antigen retrieval was performed with 0.1% Triton-X at room temperature for 10 min. Tissue sections were blocked for non-specific binding using background buster (Innovex Biosciences) at room temperature for 45 min. Sections were then incubated overnight at 4°C with primary antibodies for murine HMGB1 (Abcam, AB79823, 1:250) or the murine monocyte/macrophage markers, anti-F4/80 (Bio-rad, MCA497GA, 1:100) and anti-CD11b (Abcam AB133357, 1:1000). The next day, sections were incubated for 1 hr at room temperature with secondary fluorescent antibodies for goat anti-rabbit Alexa Fluor (AF) 488 (Abcam, AB150081, 1:500) or goat anti-rat AF595 (Abcam, AB150160, 1:500). Sections were mounted with VECTASHEILD DAPI anti-fade mounting medium (Vector, H-1200) and allowed to set for 30 min before imaging with an Axio Observer (Zeiss) using 20x/0.5 Plan-Neofluar objective, Axiocam 702 mono camera, and Zen software. Exposure settings were fixed across all tissue sections during imaging.

For quantification of protein expression fluorescent images were converted to 8-bit. A mean fluorescence intensity (MFI) was calculated from the grayscale version of the image and fluorescence channel of interest using mean grey value function in ImageJ (NIH) software. MFI measurements were taken within custom defined ROIs capturing individual IVD tissue compartments (NP, AF). ROIs were drawn blind to experimental groups.

### Cellularity measurements

Using the ImageJ software (NIH), DAPI-stained nuclei on histological sections within a custom-defined NP or AF specific ROIs were converted (8-bit), an auto-threshold was applied, and image was made binary. The cell number was then computed using analyze particles function. Cell number calculations and ROI delineation were performed blinded to experimental group and analyzed comparing multi-level pooled discs between groups. (N=3-6 per genotype per time point).

### Mechanical testing

Individual IVDs were mechanically tested on a TA Electroforce DMA 3200 Mechanical Tester. Prior to mechanical testing IVDs were thawed in PBS at 37°C for 1 hr. Immediately prior to testing, each sample was imaged using fluoroscopy to measure geometric properties using ImageJ software (NIH). Disc height was measured in mm between visible boney EPs, and cross-sectional area of the discs were approximated using the area of an ellipse (mm^2^). Unconfined compression testing between two impermeable platens where force and displacement was recorded using WinTest software. A 0.02N preload was applied to each sample followed by 20 cycles of sinusoidal loading at 0.1 Hz to a maximum load of 0.25N, corresponding to approximately 1X body weight [39, 48, 49]. Dynamic Modulus (MPa) was calculated from the ratio of the applied stress and measured strain during the 20th cycle of loading after a repeatable hysteresis pattern was established. Following dynamic loading, a creep load ramp of 0.25N was applied and maintained for 1200 seconds [39, 48, 49]. The resulting creep strain (mm/mm) was quantified at the end of the hold, representing equilibrium properties. Lastly, an equilibrium modulus (MPa) was calculated from the ratio of the equilibrium stress and strain. Mechanical testing across groups and within time points was performed on the same day. Measurements were analyzed using multi-level pooled discs between groups (N=3-6 per genotype per time point).

### Disc conditioned media – macrophage migration assay

IVDs from control and HMGB1^KO^ mice were harvested 3-days following tamoxifen IP injections (N=3 per group). Whole IVDs containing NP, AF, and EPs, were washed in sterile HBSS and cultured in complete DMEM/F-12 containing 20% FBS (Crystalgen) and 1% penicillin-streptomycin (ThermoFisher) for 24 hrs to allow for organ culture equilibrium. Following initial equilibrium culture IVDs were either left intact or punctured using a 26G needle inserted two-third of the IVD width, guided by custom cut 20G needle sleeve. Intact and punctured IVDs from; control and HMGB1^KO^ mice, IVDs were cultured in 24-well plates (three IVDs per well) within 700uL of serum free media containing DMEM/F-12 with 1% penicillin-streptomycin for 48 hrs. Following 48 hrs supernatant media from each well was collected and used as conditioned media.

Macrophages were isolated from the femurs and tibias of C57BL/6 (Jackson laboratory) mice. Bone marrow from isolated femurs was flushed out with complete RPMI media containing 20% heat inactivated FBS (Benchmark Gemini) + 1% antibiotic/antimitotic using a 27G needle and 1mL syringe. A naïve BMDM population (M0) was obtained by adherence to non-treated plastic following 7 days of culture with complete RPMI 1640 supplemented with 20% L929 conditioned media (LCM) containing macrophage colony stimulating factor (M-CSF) [50].

Following growth culture, M0 macrophages were seeded onto 8.0μm polycarbonate transwell inserts (6.5mm, Costar #3422) at 100k cells per insert, within 24-well plates. Serum free RPMI containing 1% penicillin-streptomycin was placed in the top well, while disc conditioned media collected from intact and punctured IVDs from; control and HMGB1^KO^ mice were placed within the bottom well. Following 24 hrs, transwell inserts were fixed in 4% PFA and the top of the transwell was swabbed with cotton-tipped applicators to remove non migratory cells. Inserts were then washed in PBS, and permeabilized in 0.1% Triton-X-100 for 10 minutes. The inserts were then washed in PBS again, stained with VECTASHEILD DAPI anti-fade mounting media (Vector, H-1200) and incubated for 15 minutes before imaging with an Axio Observer (Zeiss) using 20x/0.5 Plan-Neofluar objective, Axiocam 702 mono camera, and Zen software. Using ImageJ software (NIH) DAPI-stained nuclei were counted (analyze particle function) as described prior (N=3 per group).

### Statistics

Differences between groups (control vs. HMGB1^KO^) were analyzed with Student’s t-test, with Holm-Sidak post-hoc for multiple comparisons. Differences across more than two groups were analyzed with ANOVA with Tukey post-hoc test. Differences of p<0.05 are considered significant. Statistical analyses were performed using GraphPad Prism (V9.4.1).

## Results

### Validation of histological degeneration following caudal needle puncture injury

To validate severe structural degeneration following needle puncture injury, a histological analysis was performed on Safranin-O (cartilage) stained injured and adjacent to injury IVD sections [47]. Clear degenerative changes, including NP and AF tissue disruption and loss of cellularity were observed within injured discs 1-day following injury, indicating successful puncture (Fig. 1B). Looking further to 14- and 28-days post injury time points, a similar degree of severe degeneration could be observed within injured discs, as depicted by a clear loss of NP and AF cellularity, loss of cartilage staining within the NP, and loss of a clearly demarcated NP:AF border (Fig. 1B). In histological analysis of adjacent level IVDs, a primarily healthy phenotype was observed at all time points (1-, 14-, and 28-days post injury) via a highly cellular and vacuolated NP, concentric lamellae within the AF, and a distinct NP:AF border (Fig. 1B). Qualitative assessment of degenerative changes was supported with significant increases in histological scores within individual NP and AF compartments, and total scores when comparing injured IVDs to both adjacent (NP: p=0.0041, AF: p=0.0093, total: p<0.0001) and basal IVDs from non-injured mice (NP: p=0.0003, AF: p=0.0070, total: p<0.0001) (Fig. 1C). No significant changes in histological scores were seen between adjacent and basal groups (Fig. 1C). Though not included within the histological grading criteria, assessment of morphologic features of the EP was performed and guided by EP grading criteria outlined in Boos et al. [3]. EP analysis revealed no significant observable degenerative changes between injured and adjacent to injury or basal IVDs (Fig. 1B). Histomorphological analyses revealed the method of needle puncture injury produced significant degenerative structural changes to injured IVDs, however adjacent IVDs maintained a healthy phenotype. Further, the structural degeneration induced by injury primarily affected the NP and AF compartments, with no observable degenerative differences in the EP.

### Needle puncture injury increases Hmgb1 and common inflammatory cytokine gene expression within injured and adjacent IVDs

Looking at the inflammatory profile following needle puncture injury, we evaluated inflammatory mediators associated with DD, including: *Hmgb1*, *Il1b*, *Tnfa*, *Il6*, and *Nos2* [7–11, 22, 23]. *Hmgb1* was upregulated at 1- (p=0.000082), 3- (p=0.0030), and 28-days (p=0.0236) post injury in injured discs compared to IVDs from basal mice (Fig. 2A). Interestingly, *Hmgb1* expression was also upregulated at 1- (p=0.00135), 3- (p=0.0030), and 28-days (p=0.00005) post injury and downregulated at 14-days (p=0.0429) within adjacent discs compared to IVDs from basal mice (Fig. 2A). Evaluating additional inflammatory cytokines, *Il1b*, *Il6*, *Nos2*, and *Tnfa*, revealed no significant differences in injured and adjacent IVDs compared to basal at 1-, 3-, 7- or 14- days post injury (Fig. S1). However, *Nos2* was upregulated in injured discs (p=0.000352) at 28-days compared to basal (Fig. S1). Expression of *Il6* (p=0.00822) and *Tnfa* (p=0.0432) were upregulated in adjacent IVDs at 28-days post injury compared to basal (Fig. S1). Gene expression results reveal upregulation in several inflammatory mediators of DD, with *Hmgb1* upregulation being seen more consistently across time points following injury. Further, increased expression of inflammatory markers in both adjacent and injured discs, suggest local inflammatory responses to injury within non-injured discs.

**Figure 2:**
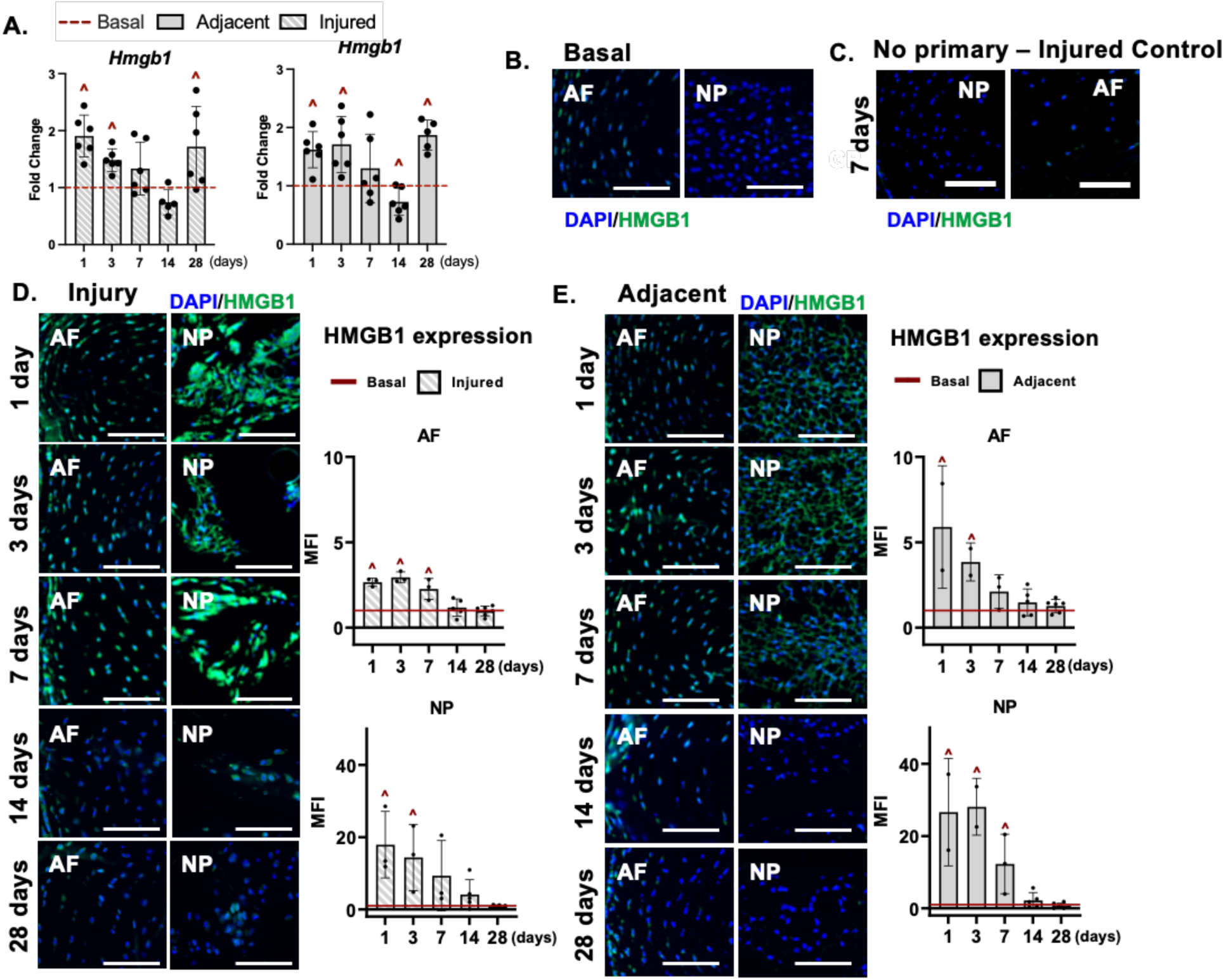
*HMGB1* activation within injured and adjacent IVDs. (**A**) Gene expression changes from total RNA isolated from adjacent and injured whole IVDs containing NP, AF, and EP, following injury (days post injury). Gene expression relative to basal whole IVDs isolated from uninjured age-matched mice. Representative images of IF staining for HMGB1 (green) within basal (**B**), injured (**D**), and adjacent (**E**) IVDs (days post injury). (**C**) Representative no primary IF control image of an injured IVD, 7 days post puncture. MFI quantification of HMGB1 expression within AF and NP compartments of injured (**D**) and adjacent (**E**) IVDs (days post injury). MFI normalized to basal IVDs from uninjured age-matched mice. Comparisons between basal and injured or adjacent discs ˄p<0.05. Scale bars = 100μm.

### Needle puncture injury produces HMGB1 protein upregulation throughout injured and adjacent IVDs

Following observations of marked increases in regional *Hmgb1* expression induced by needle puncture injury, we next evaluated HMGB1 protein expression. Beginning with injured discs, we observed clear increases in HMGB1 protein expression as indicated by increased staining intensity within the AF and NP at 1-, 3-, 7-, and 14-days, with a more mild presence at 28-days post injury when compared to a basal IVD (Fig. 2B,D). Quantification of compartment specific MFI revealed significant increases in HMGB1 protein expression at 1- (p=0.0013), 3- (p=0.000617), and 7-days post injury (p=0.0155) in the AF, and at 1- (p=0.00144) and 3-days (p=0.00558) post injury in the NP when compared to compartment specific basal MFI levels (Fig. 2D). Similarly, evaluation of HMGB1 protein expression within adjacent IVDs compared to basal revealed observable increases in HMGB1 staining intensity within the NP and AF at 1-, 3-, and 7-days post injury (Fig. 2B,E). MFI quantification in the adjacent IVDs mimicked what was observed with significant increases in HMGB1 protein expression in the AF at 1- (p=0.007) and 3-days post injury, and in the NP at 1- (p=0.00112), 3- (p=0.00001), and 7-days (p=0.00646) post injury when compared to basal MFI values (Fig. 2E). No significant differences between basal and adjacent IVDs in HMGB1 protein expression were observed at 14- and 28-days post injury. The robust increase in protein expression sustained up to 7-days post injury in both adjacent and injured IVDs compared to basal discs from uninjured mice further supported the gene expression results and provide further evidence of increased HMGB1 production in response to tissue damage.

### Validation of IVD specific Cre recombination and HMGB1 knockout in tamoxifen injected uninjured and injured mice

HMGB1 gene and protein expression within basal, uninjured, IVDs was evaluated 3-days following tamoxifen IP injections. Qualitative analysis of HMGB1 at the protein level revealed an observable decrease in HMGB1 expression throughout the AF, NP, and EP of HMGB1^KO^ discs compared to the control, as indicated by fluorescent imaging (Fig. 3A). In compartment specific gene expression analysis, a significant decrease (p=0.032) in *Hmgb1* gene expression within the NP was observed within HMGB1^KO^ mice compared to control (Fig. 3B). No significant difference in *Hmgb1* gene expression was observed within the AF across groups (Fig. 3B). This lack of significant downregulation of *Hmgb1* gene expression within AF tissue of HMGB1^KO^ mice is likely due to the mixed population of cells harvested during AF isolation, while a homogenous NP cell population contained within the NP region is more easily harvested. Regardless, validation results showed this *in vivo* mouse model targets all IVD tissue types for deletion of *Hmgb1*, resulting in decreased expression throughout the IVD.

**Figure 3:**
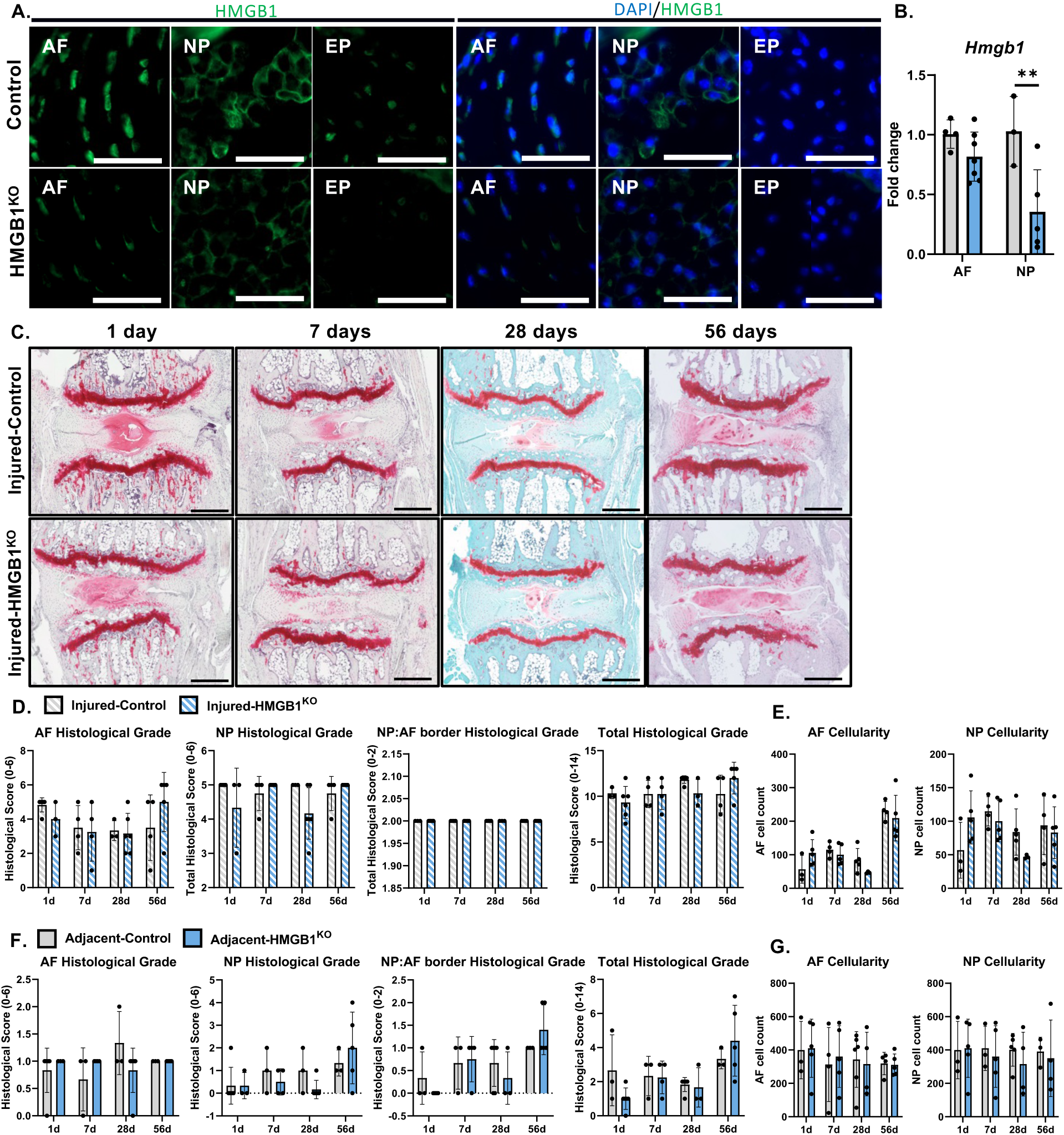
*HMGB1* knockout produces no significant protection to IVD health following injury. (**A**) Representative images of IHC staining for *HMGB1* (green) within HMGB1^KO^ and control IVDs 3-days following IP injections. Scale bar = 100μm. (**B**) Gene expression of *Hmgb1* within pooled AF or NP compartments from IVDs 3-days following IP injection. Fold change expressed relative to control within compartment. **p<0.01. (**C**) Representative images of safranin-O stained mid sagittal sections of control and HMGB1^KO^ discs 1-, 7-, 28-, and 56-days following injury. Scale bar = 500μm. (**D,F**) Histological scoring within NP, AF, and NP:AF border compartments, and Total score. (**E,G**) Quantification of AF and NP cellularity within hand drawn ROIs of DAPI nuclear stained mid sagittal sections.

### IVD HMGB1 knockout produces spontaneous caudal DD by 3-months

An initial characterization of *Hmgb1* knockout under basal (non-injured) conditions was then performed. First, structural differences were assessed across HMGB1^KO^ and control caudal IVDs, 3-months post injection, or Cre recombination. Primarily, control and HMGB1^KO^ discs appeared healthy at 1-, 2-, and 3-months post injection time points. This was characterized by highly cellular uniform NP compartments, concentric lamellae within the AF, and uniform cartilaginous EPs (Fig. S2). Interestingly, spontaneous severe DD (total histology score 8 or greater) was observed within 4/15 HMGB1^KO^ discs evaluated at 3-months post injection (Fig. S2). This was characterized by a complete loss of NP compartment cellularity and structure and a loss of cellularity and lamellae alignment, and presence of rounded chondrocyte-like cells throughout the AF (Fig. S3). Due to the sporadic nature of this severe DD phenotype within HMGB1^KO^ mice no significant differences in total histological scoring was found between HMGB1^KO^ and control mice at 1-, 2-, or 3-months post injection time points (Fig. S2). No significant changes were observed between HMGB1^KO^ and control groups in AF or NP cellularity at any time point evaluated (1-, 2-, and 3-months) (Fig. S2). Lastly, functional assessment of disc height changes revealed no significant differences between HMGB1^KO^ discs compared to control (Fig. S2). Initial structural assessment of HMGB1^KO^ revealed no significant changes across caudal disc health, as indicated via histological grading, IVD cellularity, and disc height. However, the observed spontaneous severe degenerative phenotype in HMGB1^KO^ discs indicates that HMGB1 plays an important role in maintaining tissue homeostasis.

In an evaluation of lumbar disc health following *Hmgb1* knockout, few significant changes were observed in histomorphological and fluoroscopy analysis. Lumbar IVDs from both control and HMGB1^KO^ mice at 3-months post recombination were observed to maintain healthy phenotypes, with concentric AF lamellae, uniform cellular NP compartments, and healthy cartilaginous EP (Fig. S3). In an evaluation of disc height, a significant increase was seen in HMGB1^KO^ disc height at 1-month (p=0.050), with no differences observed at 2- or 3-months post injection (Fig. S3). Ultimately, histological, cellular, and disc height analysis revealed an *Hmgb1* knockout in lumbar disc cells to produce no significant effects, protective or detrimental, to IVD health by 3-months.

### Hmgb1 knockout within IVD cells produces no significant protection to regional or injured IVD health following injury

To identify whether HMGB1 plays a primary role in maintaining tissue health following injury, histomorphological analysis of tissue health was performed on injured IVDs up to 56-days following injury. First, HMGB1^KO^ validation was performed on IVDs following needle puncture injury, as previous results supported a robust increase in HMGB1 positive staining with injury (Fig. S4). Increased HMGB1 staining was observed most notably within injured control IVDs throughout the NP at 1-, 7-, 28-, and 56-days, and throughout the AF at 28-, and 56-days post injury when compared to injured HMGB1^KO^ discs (Fig. S4). Quantification of HMGB1 staining revealed a significant decrease in HMGB1 expression within the NP of injured HMGB1^KO^ discs at 1- (p=0.0259) and 28-days (p=0.0057) post injury when compared to injured control IVDs (Fig. S4). The varied results in later time point analyses of *HMGB1* protein expression within the AF of HMGB1^KO^ mice may be influenced by mixed cell populations, not targeted by the *AcanCre^ERT2/+^*, being recruited to the IVD following injury. However, HMGB1^KO^ protein expression results show a still significant decrease in *HMGB1* expression within the NP of HMGB1^KO^ discs following injury.

In analysis of histological changes following injury, severe DD was observed across both HMGB1^KO^ and control injured discs at all time points (1-, 7-, 28-, 56-days) (Fig. 3C). Needle puncture injury produced structural changes including a complete loss of NP cellularity and structure, a loss of AF cellularity and concentric lamellae structure, and a complete loss of delineation between the AF and NP (Fig. 3C). In quantifying these changes, histological scoring reflected the observable changes with no significant differences between HMGB1^KO^ and control injured IVDs at any time point within total or compartmental categories (AF, NP, and AF:NP border) (Fig. 3C,D). No significant differences were observed in AF or NP cellularity following injury at any time point (Fig. 3E). Histological analysis suggests the severe structural degeneration induced by needle puncture is largely not influenced by *Hmgb1* knockout within IVD cells, during the onset early of degeneration post injury, and up to 56-days post injury.

Histological analysis of IVDs adjacent to injury were assessed within HMGB1^KO^ and control mice. As was seen within prior results of wild-type mice, adjacent IVDs maintained a healthy phenotype within both HMGB1^KO^ and control groups with no significant difference between HMGB1^KO^ and control adjacent IVDs in histological scoring or cellularity measurements at any time point (Fig. S5, 3F,G). Together, these results suggest that overall structural health of adjacent IVDs is not affected by the presence of *Hmgb1* knockout within IVD cells, as indicated by similarly healthy phenotypes across HMGB1^KO^ and control mice.

### Hmgb1 knockout within IVD cells produces a transient protection from disc height loss following injury

Functional analysis of disc health was next evaluated via changes to overall disc height within adjacent and injured IVDs. First, observations of fluoroscopy imaging revealed an observable increase in injured disc height within HMGB1^KO^ mice compared to control (Fig. 4A). These observations were confirmed by DHI quantification, where an increase in DHI was observed within HMGB1^KO^ injured discs at 7- (p=0.0270) and 28-days (p=0.00264) when compared to injured control discs (Fig. 4B). No significant differences were observed between control and HMGB1^KO^ adjacent IVDs (Fig. 4B). Further, compared to baseline, HMGB1^KO^ disc height was increased at all post injury time points in both injured and adjacent IVDs (Fig. 4B). Disc height results suggest that an IVD *Hmgb1* KO promotes increases in disc height, a key clinical indicator of disc health, within both adjacent and injured IVDs.

**Figure 4:**
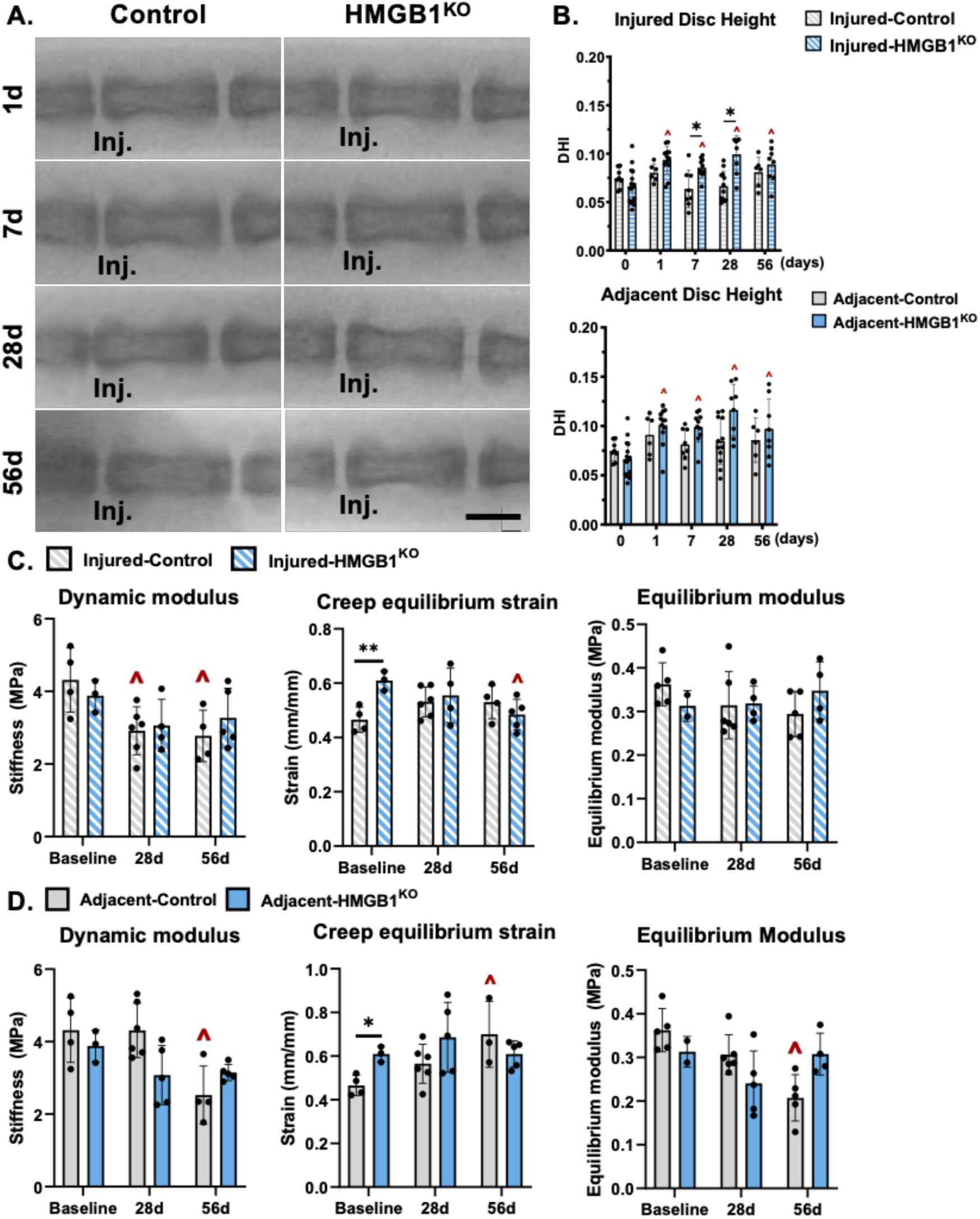
*HMGB1* knockout provides a possible protection from needle puncture injury mediated tissue compositional and mechanical property losses. (**A**) Representative fluoroscopic images of control and HMGB1^KO^ C5-C7 discs 0- (basal), 1-, 7-, 28-, and 56-days post injury. Scale bar= 1mm. (**B**) Disc height quantified via DHI of control and HMGB1^KO^ discs. *p<0.05. (**C,D**) Dynamic modulus (MPa), creep equilibrium strain (mm/mm) and equilibrium modulus (MPa) of uninjured (baseline) control and HMGB1^KO^ mice, and from injured and adjacent to injury discs 28- and 56-days post injury. Comparisons between control and HMGB1^KO^ discs *p<0.05. Comparisons within groups between baseline and post injury (ex. Baseline control vs. 56-days post injury control) ˄p<0.05.

### Hmgb1 knockout within IVD cells protects against loss of tissue mechanical property following injury

To assess the effect of HMGB1 knockout on IVD mechanics, an analysis of baseline (uninjured) control and HMGB1^KO^ discs was performed. No differences were observed in dynamic or equilibrium moduli between HMGB1^KO^ and control discs at baseline, however, a significant increase (p=0.021) in creep equilibrium strain was observed in HMGB1^KO^ vs. control (Fig. 4C). In further functional analysis, compressive mechanical properties were assessed in injured IVDs. A significant decrease in the dynamic modulus at 28- (p=0.0140) and 56-days (p=0.0135) post injury was observed in injured control discs compared to baseline (Fig. 4C). However, no significant differences between injured HMGB1^KO^ discs and baseline HMGB1^KO^ discs was seen (Fig. 4C). No differences were observed in dynamic modulus between HMGB1^KO^ and control injured discs (Fig. 4C). When comparing creep equilibrium strain between control and HMGB1^KO^ discs, no differences were observed at 28- or 56-days post injury, however, a decrease between baseline and 56-days post injury HMGB1^KO^ discs was observed (p=0.0255) (Fig. 4C). No significant differences between groups or post injury were observed within the equilibrium modulus (Fig. 4C). Given results that injury to control, but not HMGB1^KO^ IVDs, caused decreases in dynamic modulus, suggests IVD *Hmgb1* KO may play a protective role against the loss of disc compressive functionality due to injury. Interestingly a baseline increase was observed in the creep equilibrium strain within uninjured mice, suggesting induction of *Hmgb1* KO may initially modulate tissue mechanics, while decreases in creep equilibrium post injury in HMGB1^KO^ but not control mice suggest additional effects from HMGB1 on tissue mechanical properties following injury.

An evaluation of tissue mechanical properties was next applied to adjacent IVDs, both at baseline and following injury. The same baseline discs previously compared to injured HMGB1^KO^ and control discs were used for comparisons with adjacent HMGB1^KO^ and control discs. Comparing the dynamic modulus of adjacent control discs to baseline control discs, a significant decrease (p=0.0031) was seen at 56-days post injury (Fig. 4D). Conversely, a significant increase in creep equilibrium strain was observed between baseline and 56-days post injury in the control adjacent IVDs (p=0.0137), while no differences in dynamic modulus or creep equilibrium strain were observed in HMGB1^KO^ mice post injury (Fig. 4D). Lastly, evaluating equilibrium modulus changes, a significant decrease (p=0.0004) was observed in adjacent control discs vs. baseline at 56-days post injury. These results further suggest that IVD *Hmgb1* KO may protect from a loss of disc mechanical function, since discs adjacent to injury in HMGB1 KO mice were protected from alterations in the dynamic modulus, creep strain, and equilibrium modulus observed post injury in in control mice.

### Hmgb1 knockout within IVD cells provides little to no protection from IVD wide inflammatory cytokine gene upregulation following injury

HMGB1-mediated regulation of inflammation within IVDs following injury was evaluated broadly via gene expression changes of common inflammatory cytokines. Interestingly, no significant differences in *Hmgb1, Nos2*, *Il6*, or *Tnfa* gene expression was observed between control and HMGB1^KO^ injured IVDs at any time point (Fig. S6). Though contrary to our hypothesis, these results suggest, knockout of *Hmgb1* did not result in a significant effect on total IVD inflammatory gene expression at the post-injury time points evaluated within this model.

### HMGB1 mediates recruitment of macrophages to the IVD following injury

Additional investigation into possible HMGB1 mediated inflammation were focused on the chemotactic potential of HMGB1. Macrophage presence was evaluated within injured IVDs by the pan-macrophage marker, F4/80. An observable F4/80+ cell population was identified within the AF of control IVDs at 28-days post injury, and throughout the NP of injured control IVDs at 1-, 7-, and 28-days post injury (Fig. 5A). Interestingly, there was little positive staining for F4/80 within injured HMGB1^KO^ IVDs, aside from the NP at 7-days (Fig. 5A). Quantification revealed a significant decrease in F4/80 expression within the NP (p=0.039) and a trending decrease in the AF (p=0.067) of injured HMGB1^KO^ discs compared to control at 28-days post injury (Fig. 5B). Macrophage characterization was further supported qualitatively via increased staining of CD11b within the outer AF and NP of control discs at 1- and 28-days, with minimal positive staining observed within HMGB1^KO^ IVDs (Fig. 5C). Little to no F4/80 positive staining could be observed within adjacent IVDs within HMGB1^KO^ and control mice (Fig. 5D). These results suggest HMGB1 may be playing a role in recruiting macrophages (F4/80+CD11b+) to IVDs following injury.

**Figure 5:**
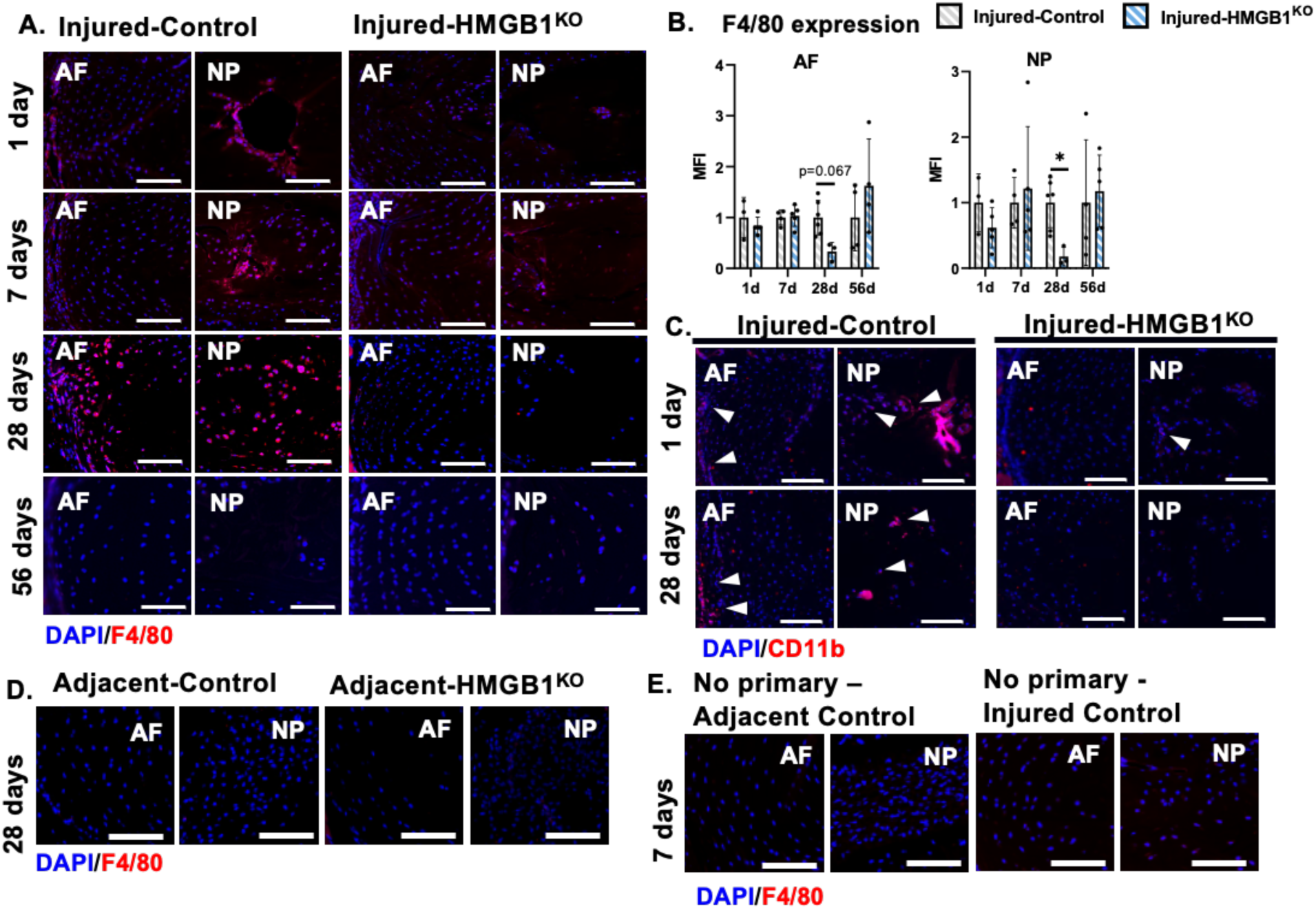
*HMGB1* knockout within IVDs following needle puncture injury. (**A**) Representative images of IF staining for F4/80 (red) in control and HMGB1^KO^ injured discs. (**B**) MFI quantification of F4/80 expression within NP and AF compartments. *p<0.05. (**C**) Representative images of IF staining for CD11b (red, white arrows) in control and HMGB1^KO^. (**D**) Representative images of IF for F4/80 (red) in control and HMGB1^KO^ adjacent discs 28-days post injury. (**E**) Representative no primary control IF images in 7-day adjacent and injured IVDs. Scale bars = 100μm.

To further confirm the potential role of HMGB1 mediating macrophage recruitment to injured IVDs, we employed an *in vitro* macrophage migration assay using intact and punctured IVD conditioned media from control and HMGB1^KO^ discs (Fig. 6A). We first observed a significant increase in macrophage migration through a transwell membrane following treatment of the cells with conditioned media from punctured control discs when compared to non-punctured disc conditioned media (p<0.0001). An injury response was also observed in HMGB1^KO^ groups, where conditioned media from punctured HMGB1^KO^ IVDs resulted in a significant increase in macrophage migration compared to conditioned media from non-punctured HMGB1^KO^ IVDs (p=0.004) (Fig. 6B,C). Interestingly, macrophage migration in conditioned media from punctured HMGB1^KO^ discs was significantly lower (p<0.0001) than punctured control discs (Fig. 6B,C). These results suggest that HMGB1^KO^ mitigates macrophage recruitment to injured IVDs (Fig. 5). The combined *in vitro* and *in vivo* findings demonstrate that HMGB1 acts as a chemokine modulator of macrophage migration in IVD puncture injury and knock out of HMGB1 mitigates its chemoattractant function.

**Figure 6:**
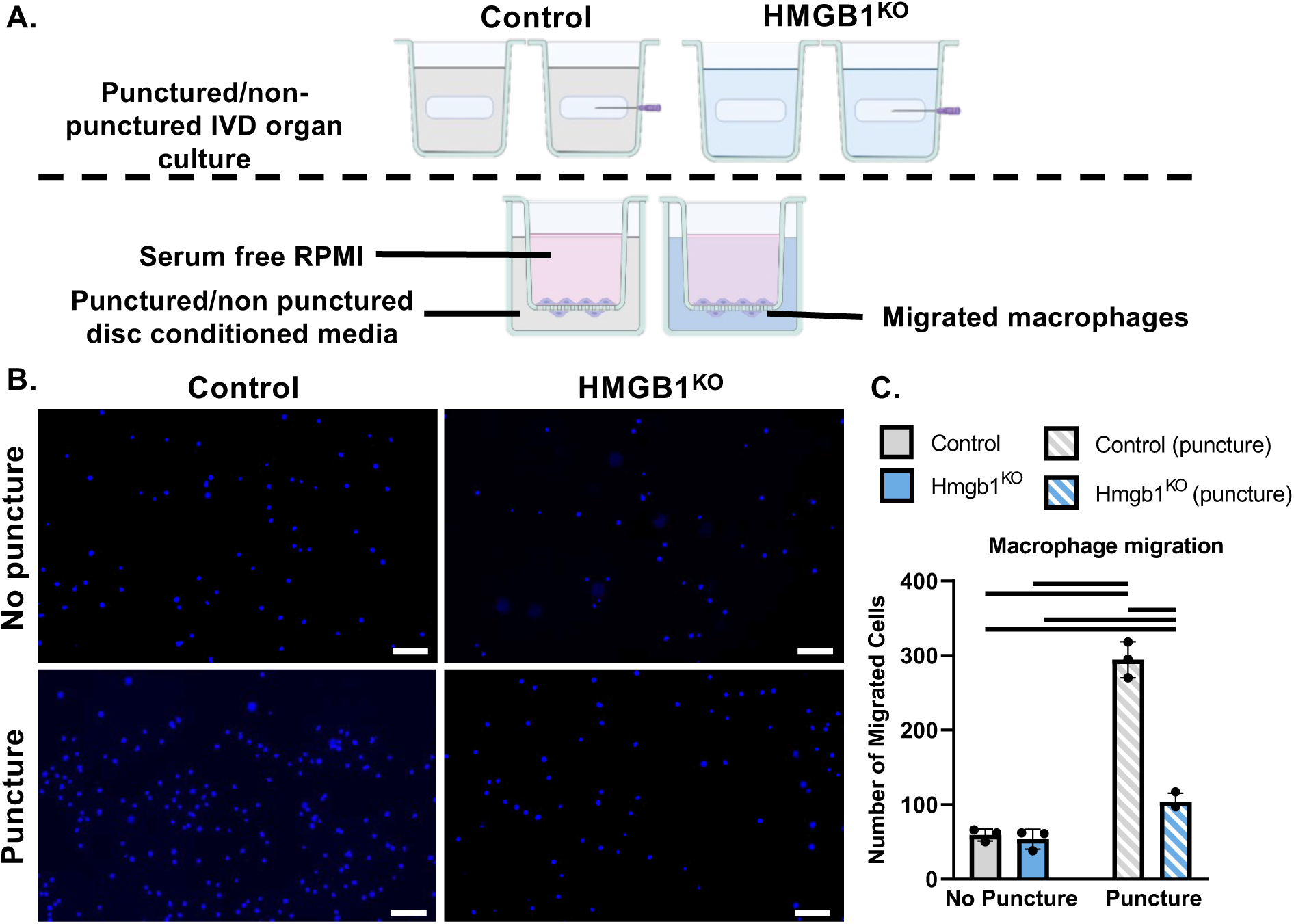
Disc conditioned media induced macrophage migration. (A) Schematic of disc conditioned media-macrophage migration assay. (B) Representative images of DAPI nuclear staining on underside of transwell insert. Scale bar 100μm. (C) Quantification of DAPI nuclear count following 24 hr stimulation with disc conditioned media from non punctured and punctured control and HMGB1^KO^ discs. Bars represent significant differences (p<0.05) between groups.

## Discussion

HMGB1 has been identified as a potent DAMP, where it is thought to influence inflammation and tissue healing during many musculoskeletal diseases [21]. The primary goal of this study was to evaluate the inflammatory roles of HMGB1 following IVD injury and their effect on tissue health, using a murine caudal needle puncture injury model. The results of this study suggest a complex role of HMGB1 following injury within the IVD, with evidence for protection from injury induced loss of tissue mechanical properties, and a transient protection from disc height losses following injury observed in HMGB1^KO^ mice compared to control. Alternatively, HMGB1^KO^ discs displayed no protection from structural degeneration or from upregulation of inflammatory markers following injury when compared to control IVDs. Further, the effects on tissue mechanics and disc height were observed consistently in the severely degenerated injured IVDs and in healthy IVDs adjacent to injury, suggesting HMGB1 is carrying out tissue maintenance functions regionally following disc injury. In evaluating possible mechanisms in which HMGB1 is responding to tissue damage, results suggest that HMGB1 is mediating macrophage recruitment to injured IVDs. Lastly, no overt changes to tissue structure were observed basally within uninjured HMGB1^KO^ mice. However, spontaneous severe DD was observed within healthy caudal IVDs from uninjured HMGB1^KO^ mice. This spontaneous severe DD was not observed in healthy lumbar IVDs. Ultimately, these novel findings provide support for HMGB1 in serving a primary role in IVD tissue maintenance, both basally and following tissue damage.

In addition to the structural and biological changes observed within IVDs following injury, we also observed a similar inflammatory profile across injured and adjacent IVDs at 28-days post injury. Utilizing IVDs harvested from basal (naïve uninjured) mice, we observed upregulation of the same inflammatory mediators, *Tnfa*, *Il6*, *Il1b*, and *Nos2* within both injured and adjacent to injury IVDs. Interestingly, opposed to biological changes, the degenerative structural change was not similar across injured and adjacent discs, with adjacent discs maintaining a healthy phenotype. Adjacent segment disease (ASD) has been previously observed clinically and in animal models in response to spinal fusion [51–54]. However, observations of ASD in response to disc puncture injury, specifically disc structural and biological changes, have not been evaluated. The results of our study provide novel findings on adjacent inflammatory profiles and rationale for investigating DD driven regional inflammation, as adjacent injury may contribute to chronic elevation of inflammatory markers, thought to be a contributor to human DD.

Part of this study evaluated the relevance of HMGB1 as a marker of disc degeneration. The results in this study, where increased expression of HMGB1 was observed in injured discs and in discs adjacent to injury, are consistent with prior findings that HMGB1 levels increase with severity in degenerated human IVDs [22, 23]. Within the puncture injury environment evaluated in this study, results reveal HMGB1 to be upregulated across the IVD, suggesting a response to initial tissue damage and a persistent disrupted microenvironment. While HMGB1 has been evaluated within a number of *in vivo* animal models of musculoskeletal disease, including bone and muscle injury, tendinopathy, OA, and RA [28–35], to our knowledge this study is the first to report on the direct effects of HMGB1 in IVD injury in vivo.

Acting as a nuclear architecture proteins, HMGB1 mediates a host of cellular functions by binding to DNA and facilitating transcription [55]. HMGB1 has also been studied for roles outside the nucleus where its functionality is dependent on redox states. Following extracellular release HMGB1 exists in a fully reduced state (fr-HMGB1), but dependent on the oxidative environment and interactions with reactive oxygen species, HMGB1 becomes oxidized into disulfide-HMGB1 (ds-HMGB1) and/or fully oxidized, ox-HMGB1. It is presently thought that fr-HMGB1 plays a chemoattractant role, through CXC motif ligand (CXCL12) and CXC motif chemokine receptor type 4 (CXCR4) interactions [56]. While ds-HMGB1 is thought to activate inflammatory responses via TLR4 binding [57], and ox-HMGB1 to mediate inflammatory resolution, though these mechanisms are still being evaluated [58]. Our results identify many possible roles for HMGB1 in mediating tissue maintenance, remodeling, and recruiting immune cells within this model. Future work will aim to elucidate how basal and injured disc environments may be influencing the extracellular HMGB1 oxidative state, and how these further influences disc health.

Prior studies evaluating HMGB1 stimulation on human IVD cells have produced mixed inflammatory effects, possibly dependent on cell type (NP, AF) and on level of degeneration severity. The pro-inflammatory effects of HMGB1 have been observed by Shah et al., with an upregulation of IL-6 and MMP-1 following HMGB1 stimulation on human NP cells [23]. Further, Fang et al., saw a similar pro-inflammatory response following HMGB1 stimulation of AF cells, through the promotion of inflammatory cytokine release, including PGE2, TNF-α, IL-6 and IL-8 [36]. Alternatively, Klawitter et al. saw HMGB1 stimulation of human NP cells from an already degenerated IVD to have an opposite effect, where decreased IL-6 expression was observed [37]. Specifically looking at the inflammatory response within the current *in vivo* study, little significant differences in inflammatory activation was observed as indicated by gene expression changes within HMGB1^KO^ mice. A possible explanation for this may be size limitations of murine IVDs requiring the analysis of gene expression across whole IVDs, not separated by region. Thus, without substantial sample sizes and pooling of multiple IVD segments, it was not possible to examine gene expression changes across individual IVD compartments within this injury study (NP, AF, and EP), providing a limitation of this study and a possible explanation for the lack of significant findings in inflammatory gene expression changes. Further evaluation utilizing methods in single cell transcriptomics would better allow for a high-dimensional characterization of inflammatory regulation dependent on HMGB1 following IVD injury.

Within the scope of inflammatory driven musculoskeletal disease, specifically arthritis, treatment with exogenous HMGB1 resulted in an increased pro-inflammatory environment as indicated by angiogenesis and enhanced recruitment of immune cells and resulting in more severe arthritic histological pathogenesis [21, 31, 32]. Further, delivery of HMGB1 into healthy murine joints resulted in inflammatory joint arthritis via activation of local macrophages and NF-κB pathway signaling [21, 31, 32]. In additional studies blocking HMGB1 within arthritic tissue resulted in an improvement in clinical arthritis scoring severity, reduced inflammation, and a reduction in migrating cells [21, 29, 34]. In specific, when HMGB1 was inhibited within LPS driven arthritic murine models a significant reduction of peripheral blood inflammatory cytokines (IL-1β, IL-6, and TNF-α) was observed [29]. Additionally, within collagen-induced arthritic mouse and rat models therapeutic HMGB1 inhibition also lead to a reduced mean arthritis score, as indicated by histological assessment [34]. Ultimately, arthritic inflammation studies largely suggest HMGB1 to worsen disease pathology. Though chronic inflammation is also commonly associated with severe DD, far less is known about whether HMGB1 exhibits a role, and whether it is harmful or protective of disease progression within the IVD. Interestingly, a mitigation of inflammatory gene expression changes following injury was not seen within this model, however we did observe protection of disc integrity following injury within HMGB1^KO^ mice, as seen through protections from disc height and mechanical property losses. These results, and the results of others, support HMGB1 to play a role in driving disease pathology, specifically tissue degradation, possibly through a regulation of local immune cell mediated inflammation, though more work is needed to identify an inflammatory regulatory role within the IVD.

Within the understood role of initiating inflammatory responses following tissue damage/injury, HMGB1 is known to act as a potent chemoattractant, recruiting mononuclear immune cells, including macrophages, fibroblasts, and neutrophils [56, 59, 60]. Specifically in the context of inflamed musculoskeletal tissues, in the previously discussed study by Pullerits et al., the intraarticular injection of HMGB1 into the murine joint space which resulted in arthritis pathogenesis was also hallmarked by a recruited cell population comprised primarily of macrophages [32]. In evaluating the chemoattractant potential of HMGB1 following injury, we observed a mitigation of macrophage recruitment to injured IVDs within HMGB1^KO^ mice, as compared to a robust recruitment of macrophages to injured control discs. This was further supported via *in vitro* migration analysis, revealing a significant mitigation of macrophage migratory function when cells were stimulated with the secretome of injured HMGB1^KO^ IVDs compared to injured control IVDs. These results demonstrate that HMGB1 participates in recruitment of macrophages following musculoskeletal tissue injury and inflammation, possibly via known mechanisms driven by the fully reduced form of HMGB1 (e.g. CXCL12/CXCR4) [61]. Regulation of macrophage recruitment could provide reasoning for the tissue scale changes in both tissue content and mechanical properties, as macrophages are well understood to mediate tissue healing and repair following injury. Future studies manipulating macrophage infiltration within this model will aim to identify whether this HMGB1 mediated immune response is a primary contributor to the observed structural and mechanical changes. Further, as macrophage infiltration was not observed in the adjacent to injury IVDs, these future studies will also aim to identify if the observed protection from adjacent level mechanical changes is driven by macrophage infiltration and degeneration within nearby injured discs. Possibly influenced by mechanisms associated with ASD, such as altered mechanical loading environment produced by neighboring mechanically compromised degenerated discs.

Though the focus of this study was to evaluate HMGB1 as a DAMP in response to IVD injury, we observed a possible homeostatic tissue maintenance role in non-injured HMGB1^KO^ mice. Following 3-months of *Hmgb1* depletion within IVD cells we observed spontaneous severe caudal DD, as indicated by histological analysis. In addition to the inflammatory and tissue healing mechanisms of extracellular HMGB1, intracellular HMGB1 plays a number of roles, including; transcriptional regulation, DNA replication and repair, telomere maintenance, and nucleosome assembly [62]. Further, HMGB1 has been identified to inhibit apoptosis and enhance cellular senescence within cancer cells [62–64]. However, HMGB1 mediated DNA repair and maintenance mechanisms and apoptotic and senescence activating functions within musculoskeletal resident cells have not been specifically evaluated. To our understanding the spontaneous severe caudal DD observed within this study is the first to suggest HMGB1 is necessary to maintain IVD health in the uninjured setting, possibly through some of its non-inflammatory related mechanisms. Ultimately these findings also present a limitation to the model system, as it remains unknown the mechanisms driving this severe DD phenotype within HMGB1^KO^ discs. Further, this degeneration was observed both in uninjured animals, as well as in adjacent to injury IVDs of HMGB1^KO^ mice, thus introducing difficulty in separating degenerative effects due to injury or due to HMGB1 depletion within IVD cells alone.

Ultimately results of this study support the complex role of HMGB1 in regard to basal disc tissue homeostasis as well as inflammatory and tissue repair mechanisms following injury. This study identified HMGB1 as a robustly activated inflammatory molecule following injury and at later stages of degeneration. Further, the identified inflammatory profile upregulated in both adjacent and injured discs suggests DD to have cascading regional effects, causing structural changes and inflammatory activation. Results suggest HMGB1 may be required for IVD tissue maintenance, as indicated by severe DD just 3-months following *Hmgb1* depletion from IVD cells. In response to injury, *Hmgb1* depletion contributed to a maintenance of disc mechanical properties and height following injury, suggestion depletion of *Hmgb1* to be beneficial. Lastly, we observed HMGB1 to play a role in macrophage recruitment following injury, providing a possible mechanism in which HMGB1 influences inflammatory and tissue healing responses within the IVD.

## Author contributions

KGB, MKK, and NOC provided study design. KGB, MKK, DCV, JRG, and GFM conducted the study. KGB, MKK, and NOC analyzed and/or interpreted the data. KGB, and NOC drafted and edited the manuscript. All authors have read and approved the final manuscript.

## Data availability statement

The data that support the findings of this study are available in the Materials and Methods, Results, and/or Supplemental Material of this article.

---

**Figure S1:**
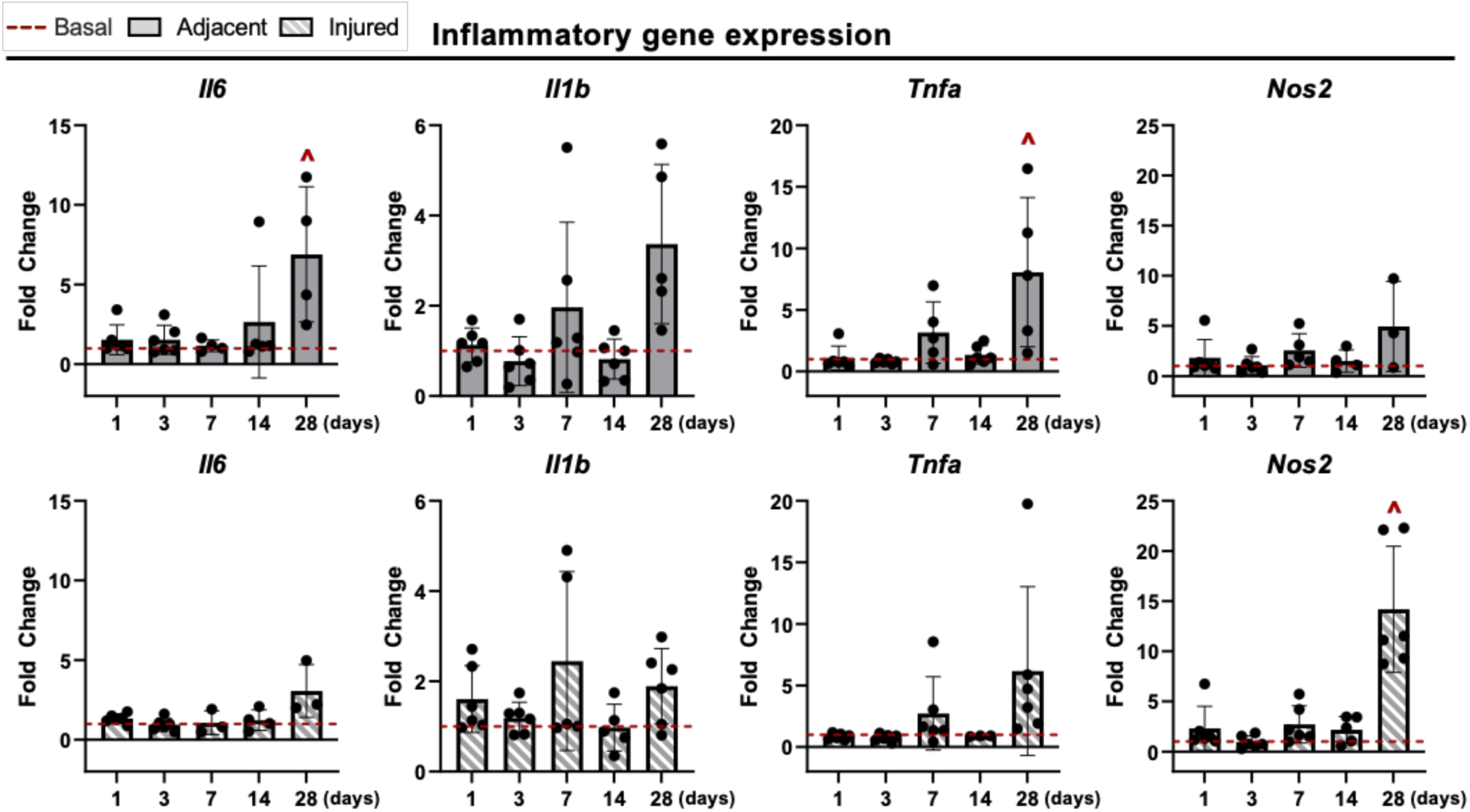
Regional inflammatory cytokine profile following needle puncture injury. Gene expression changes from total RNA isolated from adjacent and injured whole IVDs containing NP, AF, and EP, following injury (1-, 3-, 7-, 14-, 28-days). Gene expression relative to basal whole IVDs (red dashed line) isolated from uninjured age-matched mice. Comparisons between basal and adjacent, and basal and injured discs ˄p<0.05.

**Figure S2:**
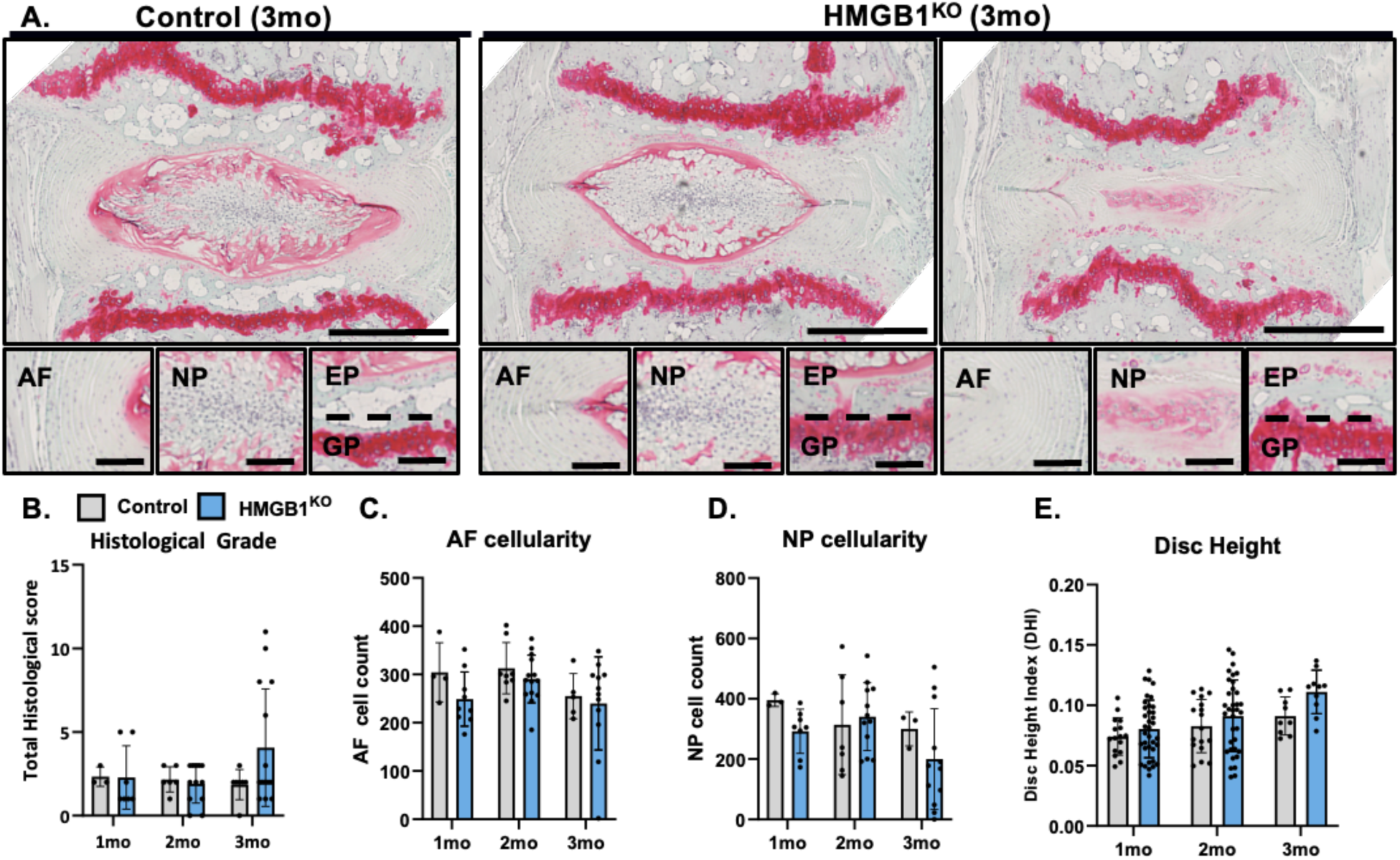
IVD *HMGB1* knockout produces spontaneous DD under basal conditions by 3- months. **(A**) Representative images of safranin-O stained mid sagittal sections of control and HMGB1^KO^ caudal discs 3-months post recombination. Scale bar = 500μm. Compartment specific zoom, scale bar = 100 μm. (**B**) Total histological scoring ranging from 0 (healthy) to 14 (most severe) of control and HMGB1^KO^ discs 1-, 2-, and 3-months post IP injection. Quantification of AF (**C**) and NP (**D**) cellularity within hand drawn ROIs of DAPI nuclear stained mid sagittal sections. (**E**) Disc height quantified via DHI of control and HMGB1^KO^ discs 1-, 2-, and 3-months post recombination.

**Figure S3:**
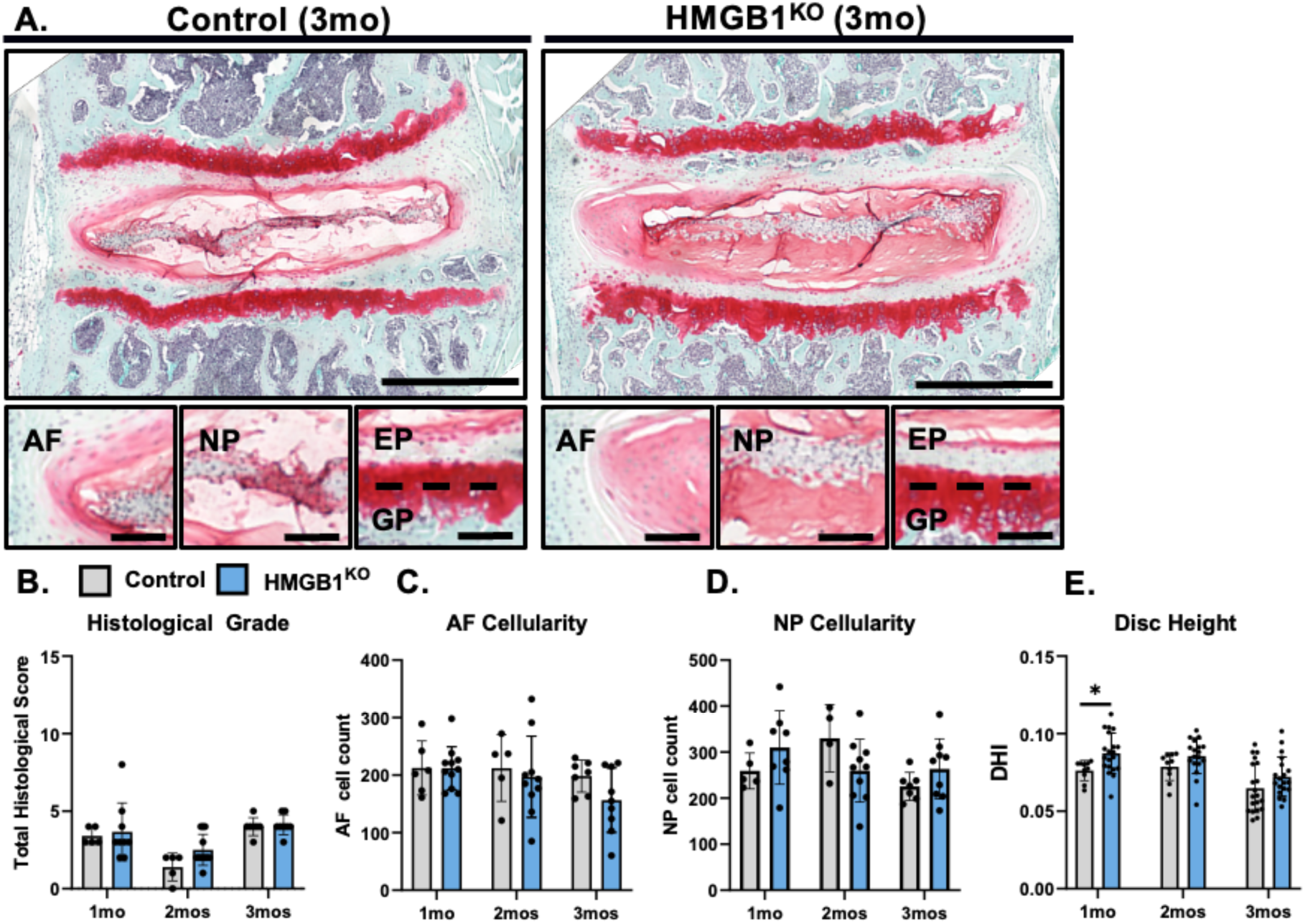
IVD *HMGB1* knockout produces no changes to lumbar disc health by 3-months. (**A**) Representative images of safranin-O stained mid sagittal sections of control and HMGB1^KO^ lumbar discs 3-months post recombination. Scale bar = 500μm. Compartment specific zoom, scale bar = 100 μm. (**B**) Total histological scoring ranging from 0 (healthy) to 14 (most severe) of control and HMGB1^KO^ discs 1-, 2-, and 3-months post IP injection. Quantification of AF (**C**) and NP (**D**) cellularity within hand drawn ROIs of DAPI nuclear stained mid sagittal sections. (**E**) Disc height quantified via DHI of control and HMGB1^KO^ discs 1-, 2-, and 3-months post recombination. *p<0.05.

**Figure S4:**
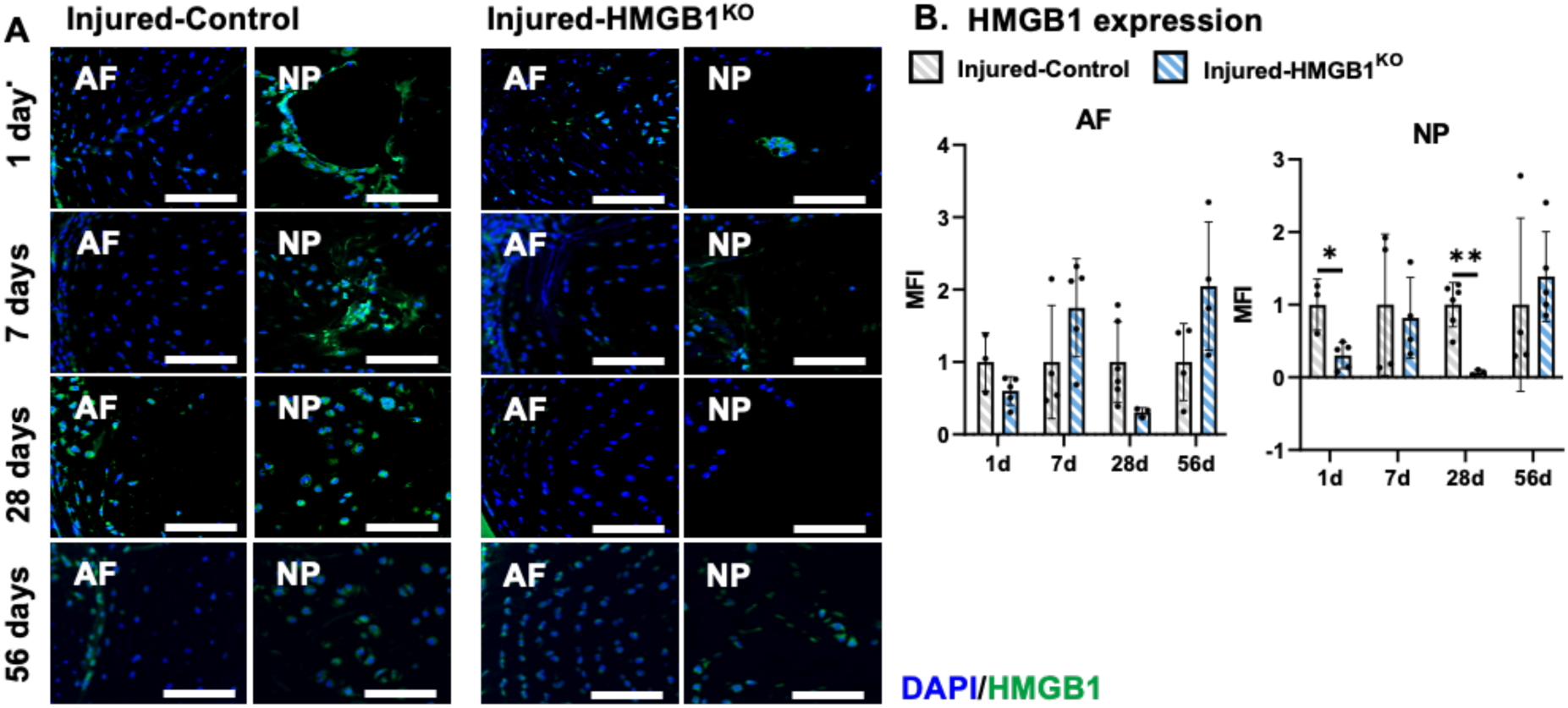
*HMGB1* knockout within IVDs following needle puncture injury. (**A**) Representative images of IF staining for *HMGB1* (green) in mid sagittal sections of control and HMGB1^KO^ caudal discs 1-, 7-, 28-, and 56-days post injury. Scale bar 100μm. (**B**) MFI quantification of *HMGB1* expression normalized to within time-point control group across NP and AF compartments. *p<0.05, **p<0.01.

**Figure S5:**
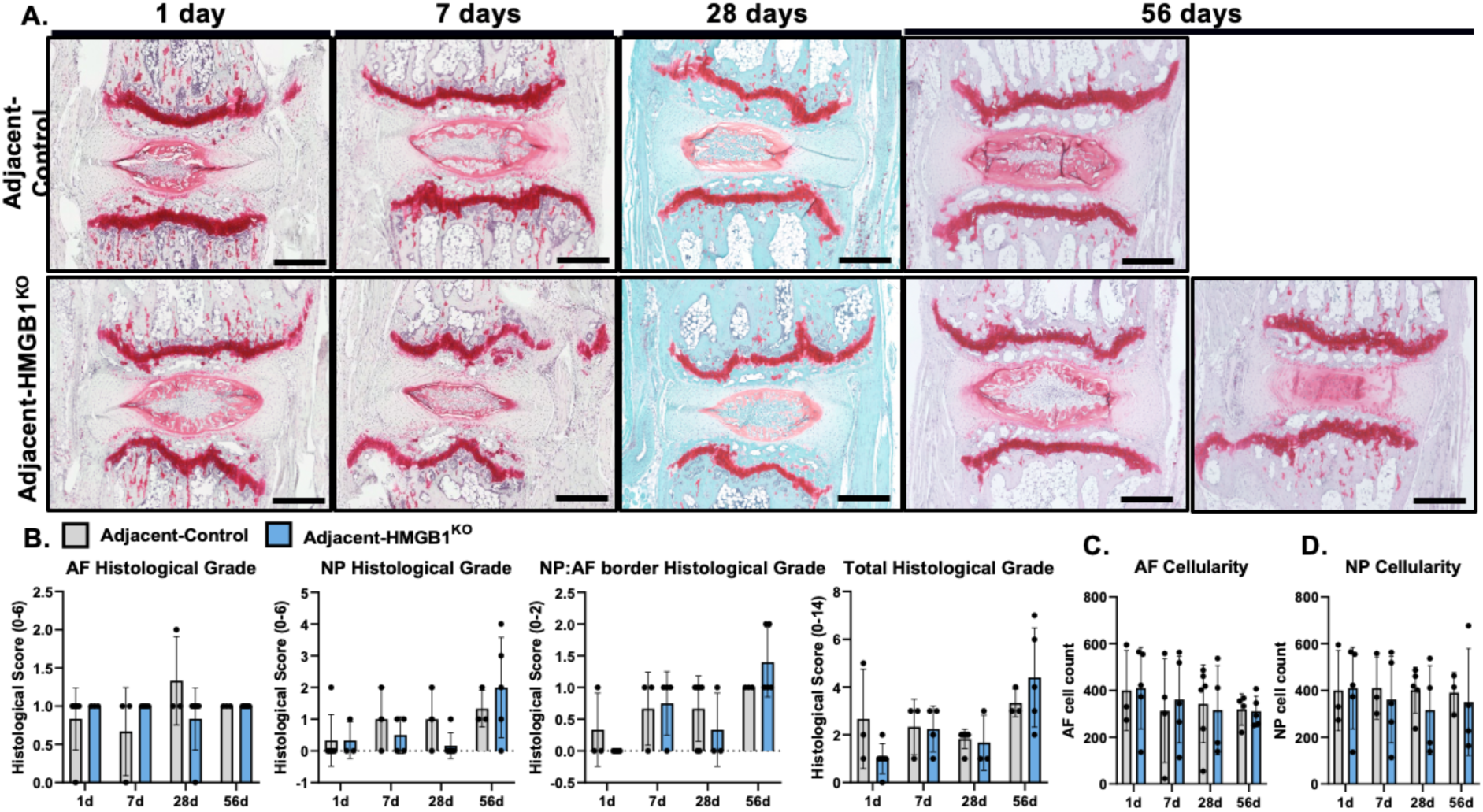
*HMGB1* knockout produces no significant protection to adjacent to injury IVD health. (**A**) Representative images of safranin-O stained mid sagittal sections of control and HMGB1^KO^ caudal discs 1-, 7-, 28-, and 56-days following injury. Scale bar = 500μm. (**B**) Histological scoring within NP, AF, and NP:AF border compartments, and Total score. Quantification of AF (**C**) and NP (**D**) cellularity within hand drawn ROIs of DAPI nuclear stained mid sagittal sections.

**Figure S6:**
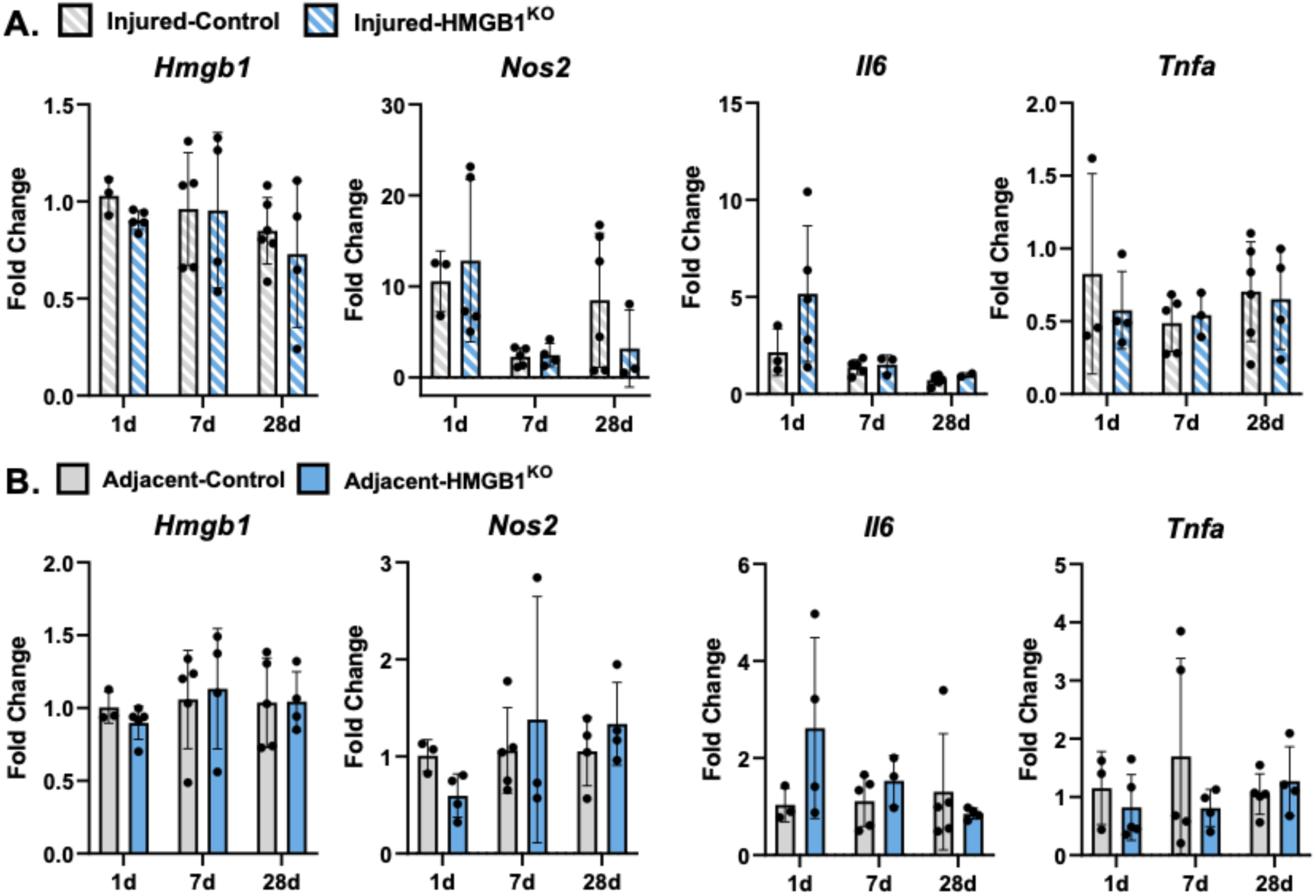
*HMGB1* knockout produced little changes to inflammatory cytokine expression following injury. (**A,B**) Gene expression changes from total RNA isolated from control and HMGB1^KO^ whole injured and adjacent to injury caudal IVDs, normalized to adjacent control IVDs.

